# Quantitative Signatures of Disassembly Mechanisms Modulating Filament and Bundle Assembly in a Shared Pool

**DOI:** 10.1101/2025.05.24.655867

**Authors:** Md Sorique Aziz Momin, Michaela Cohen, Lishibanya Mohapatra

## Abstract

How cytoskeletal structures control their assembly while sharing a common pool of their constituent parts is an open question in biology. Experiments indicate that mechanisms promoting the disassembly of these structures and replenishing the pool may play a vital role. Here, we compare the role of two modes of disassembly: monomer loss and loss of fragments (severing), in the assembly of bare filaments and bundles, modeled as a collection of linear filaments. Using analytical calculations and simulations, we show that severing can accelerate the assembly of these structures and ensure their precise size control in a shared pool. We also examine their length fluctuations and find that the decay in the autocorrelation function is faster with severing. Our study identifies parameters that influence assembly kinetics as well as the decay in autocorrelations of length fluctuations. These findings provide a framework for designing experiments that can differentiate between size control mechanisms in cytoskeletal structures.

## INTRODUCTION

Cytoskeleton dynamics play a central role in regulating various aspects of cellular behaviour, including cell morphology, motility, and intracellular transport. To perform these functions, cytoskeletal monomeric proteins such as tubulin dimers and actin monomers often assemble into higher-order structures known as bundles [1–4]. These bundles exhibit different morphologies related to their functions. For example, uniformly sized actin bundles called microvilli (typically a few microns in length) line the small intestines and are crucial for enhancing cellular adhesion and nutrient absorption [5–7]. Stereocilia, also composed of actin filaments (ranging from 1 to 120 microns in length), display a staircase-like structure and play an essential role in signal transduction in the inner ear [8, 9]. An example of a microtubule-based bundle is cilia, which are used for motility and sensory functions in various cell types [10, 11]. Precise assembly of these structures is essential—microvilli defects are linked to celiac disease [12, 13], stereocilia dysfunction causes hearing and balance disorders [14], and ciliary defects underlie primary ciliary dyskinesia and polycystic kidney disease [15, 16].

In many cases, the cell regulates the sizes of these structures, as seen in the case of unicellular algae *Chlamydomonas reinhardtii*, which uses two equal-sized flagella for locomotion. When one flagellum is severed, the cut flagellum regrows, while the intact one shortens to ensure both reach an equal length [17, 18]. Additionally, bundles display a range of assembly times. In budding yeast, actin cables form within seconds to minutes at elongation rates of 0.3–0.5 µm/s [19–22]. Microvilli in frog epithelium assemble in minutes [5], neuronal filopodia elongate over tens of minutes [23], and *Chlamydomonas* flagella require hours [17]. In contrast, stereocilia in mouse inner hair cells take several days to form [24]. Remarkably, these structures assemble from a pool of monomeric proteins (tubulin or actin monomers) while undergoing a rapid turnover of their internal components [25–34], and are still able to assemble to their specific size in time. Mechanisms that control the assembly of these structures in a shared pool of their components have thus been an active area of research in cell biology. Experiments have revealed that a variety of proteins help shape the size and organization of these cytoskeletal structures by affecting their assembly or disassembly [26, 35]. In particular, disassembly factors [36, 37] have been observed to control the size of various organelles [34, 36, 38, 39], such as kinesin-13 in the flagella of *Giardia* and katanin in neurons [36, 37], as well as mechanisms that modulate assembly like intraflagellar transport (IFT) [25, 26, 40].

Disassembly mechanisms can play an important role in remodeling cytoskeletal structures. In addition to dismantling pre-existing structures and replenishing the pool of building blocks, these mechanisms can also provide crucial length-sensing feedback to regulate the size of individual structures sharing a common pool (e.g., via depolymerizing kinesins such as KIP3 and severing proteins [41–44]). Several studies have shown that when multiple structures compete for a shared subunit pool, negative feedback between growth rate and size can ensure robust size control [32, 34, 45]. However, in the case of bundles, Rosario et al. [46] challenged the need for filament-level feedback for bundle length regulation. Using a theoretical model, they demonstrated that when filaments are crosslinked into bundles, the geometric constraints alone can lead to emergent length regulation (without invoking explicit feedback mechanisms). This raises further questions: How might the bundle architecture and other size-dependent feedbacks influence bundle assembly kinetics and/or their length fluctuations? Intriguingly, in the case of actin cables in budding yeast, McInally et al. [47] combined quantitative imaging with theoretical modeling to show that the spatial arrangement and turnover of filaments within actin cables—rather than individual filament dynamics—can explain observed assembly kinetics. This finding highlights the role specific disassembly mechanisms (like filament turnover in cables) can play in modulating the assembly kinetics of bundles. For these reasons, in this study, we explore the effect that specific disassembly mechanisms (with and without negative feedback at the individual filament level) can have on the assembly kinetics of filaments and bundles assembling within a shared pool environment.

Disassembly can manifest in many ways for a cytoskeletal structure. A structure can lose one monomer at a time or lose portions of filaments, as observed in the case of severing. There is evidence that severing mechanisms play a role in modulating the size of cellular structures in vivo, such as microtubule lengths in neurons [36, 48] and spindle size [43, 49, 50]. More recently, it has been shown that severing can play a critical role in the stress relaxation behavior of dynamic actin filament networks by influencing their kinetic rates of assembly, disassembly, and filament length [51]. Severing of filaments can arise from various mechanisms, including tensile forces [52], externally applied forces like thermal stress [53], filament bending and buckling [54, 55], and also from specific severing proteins. These proteins operate by cutting protein polymers at their binding sites; longer polymers, with their increased number of binding sites, are more likely to be severed, indicating a length-dependent disassembly rate [56, 57]. While numerous theoretical and experimental studies have investigated the properties of the severing mechanism to control size [44, 46, 57–61] on various cytoskeletal structures, their precise impact on the assembly kinetics and length fluctuations of filaments and bundles within a shared pool environment has not been explored.

Using analytical calculations and stochastic simulations, we model the assembly of two types of cytoskeletal structures (bare filaments and bundles) in a shared pool, which undergo disassembly by losing either individual monomers or portions of filaments. We begin by characterizing the assembly of individual filaments and bundles and later investigate their simultaneous assembly in a shared pool. In each case, we examine how the assembly kinetics, steady-state size distribution, and size fluctuations vary over time at steady state. A key goal is to distinguish the effects of the two disassembly mechanisms—losing monomers or portions of filaments—on each structure in different situations. We identify parameters that describe both the assembly kinetics and the decay in autocorrelations of length fluctuations. These findings provide a framework for designing experiments to better understand the role of disassembly mechanisms in cytoskeletal structure formation and maintenance.

## RESULTS

### Assembly of Individual Filaments in a Pool

We first examine the assembly of bare filaments in a pool of their monomers. The addition rate is proportional to the monomer pool, which may be either limited or in excess (free pool). Filaments disassemble via monomer dissociation from the ends, replenishing the pool, or by losing portions of the filament, with the severed part of the filament becoming part of the pool. We compare filament assembly in scenarios where filament length is controlled: assembly with constant disassembly in a limited pool, assembly with severing in a free pool, and assembly with severing in a limited pool. While filament assembly with constant disassembly in a limited pool has been studied previously [32], we reproduce these results to highlight the impact of severing on filament kinetics and steady-state length distributions. Since the results for severing in a free pool and severing in a limited pool are similar under our chosen parameters, they are presented in the Supplementary Material, while the main text focuses on the constant disassembly in a limited pool and severing in a free pool mechanisms (see Figures 1A–1B).

**Figure 1.**
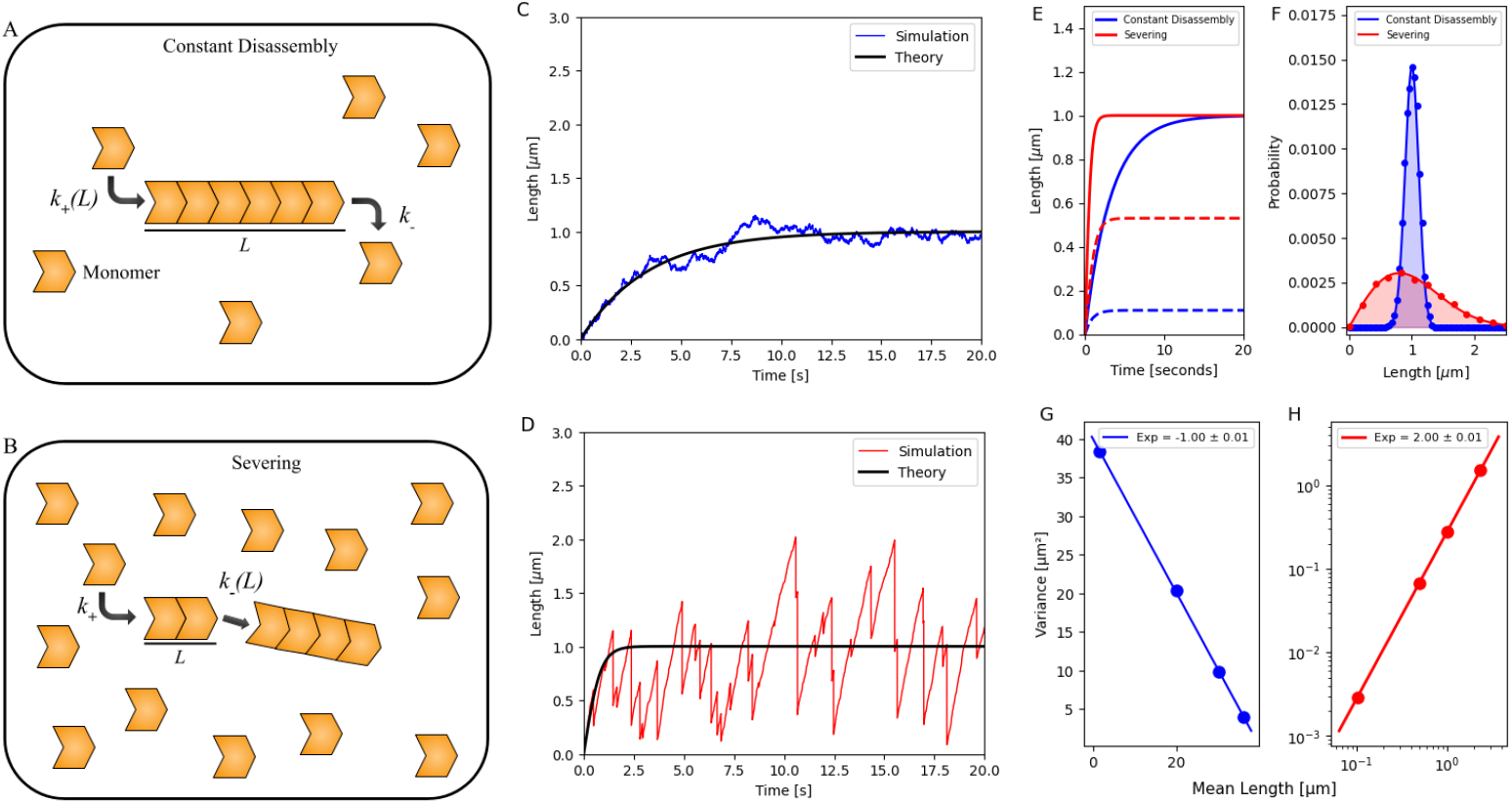
Growth dynamics of a bare filament regulated by (1) constant disassembly in a limited pool and (2) severing in a free pool. (A, B) Schematics showing the growth of a bare filament regulated by (1) constant disassembly in a limited pool and (2) severing in a free pool. (C, D) Stochastic simulation of the growth trajectory of a bare filament regulated by (1) constant disassembly in a limited pool (blue) and (2) severing in a free pool (red). The simulations are overlaid with the analytically obtained results (black) in the Supplementary Material. (A) Mean length (solid line) and standard deviation (dashed line) of bare filament calculated from simulations for the two mechanisms, plotted versus time. (B) Probability distribution of the steady-state length of bare filaments regulated by the two mechanisms. Dots denote data obtained by simulation results, while solid lines represent analytical prediction (as detailed in the Supplementary Material). (G, H) Variance of filament length distributions is plotted against the mean filament lengths regulated by the two cases. A normal scale was chosen for the constant disassembly mechanism and a log-log scale for the severing mechanism. For panels (C–F), parameters used *N* = 1000 monomers (each 4 nm in size), with 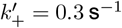, *k*_−_ = 225 s^−1^ (constant disassembly), and *s* = 0.0075 monomers^−1^ s^−1^ (severing). For pan-els (G–H), *N* = 10000, and different 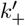 values were used to achieve distinct mean filament lengths. In the constant disassembly scenario: 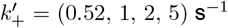 and *k*_−_ = 5000 s^−1^; in the severing scenario: 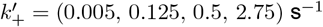 and *s* = 0.125 monomers^−1^ s^−1^.

*Assembly with constant disassembly in a limited pool:* We consider the assembly of a filament in a pool where the total number of monomers, *N*, is fixed, and monomers associate with the filament at a rate proportional to the free monomers (*N* − *L*), where *L* is the filament length. The growth rate is hence 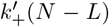, where 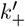 is derived by dividing the second-order rate constant for monomer addition by the cytoplasmic volume. The assembly rate decreases as the filament grows, while the dissociation rate remains constant at *k*_−_. At steady state, these rates balance, leading to a peaked filament length distribution, as confirmed by stochastic simulations [62, 63]. The evolution of mean filament length ⟨*L*⟩ can be described by the equation:

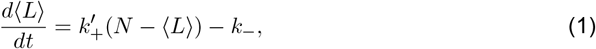

the solution to this equation is given by 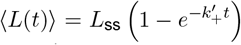, where 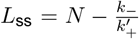 repre-sents the steady-state filament length, and 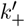 determines the relaxation rate to steady state (Figure 1C).

*Assembly with severing in a free pool:* In this mechanism, we assume that filaments assemble at a constant rate proportional to the monomers, *N*, but can disassemble by losing portions of the filament (i.e., get severed). Monomers on the filament are chosen at random (with a rate *s*), and once the site is chosen, monomers to the right of the chosen site on the filament (*L*_*f*_) are lost and reincorporated into the pool. The choice of monomer site where the portion of the filament gets cut scales with filament length, and hence longer filaments will likely disassemble more. Concomitantly, the disassembly rate (i.e., rate of subunit loss) is also length-dependent, but scales quadratically with length, ≈ *s*⟨*L*⟩*L*_*f*_ . The evolution of the equation for mean length can then be described by the following equation:

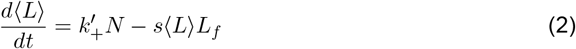

where the rate of growth is 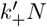. Just like in the previous case, this equation predicts that, after a growth phase, the filament would reach a steady state when the assembly rate matches the severing rate. This is confirmed in our stochastic simulations [62, 63], where we also observe large fluctuations in length (Figure 1D), which pertain to events where portions of filaments are lost.

In general, 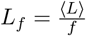, and the steady-state solution for Equation 2 is 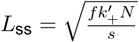. Compar-ing this expression with the exact solution to the master equations for this process, previously solved in [51, 61, 64], we find that 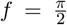. Using this, Equation 2 can be solved to obtain ⟨*L*(*t*)⟩ = *L*_ss_ tanh(*k*_*S*_*t*), where 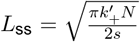 and 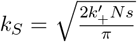, determining the rate at which steady-state length is achieved. This, along with a trajectory obtained from stochastic simulation, is plotted in Figure 1D.

In the Supplementary Material, we also consider the growth of filaments (that can sever) in a pool that is limited. We find that, for the choice of our parameters, the simulation results are similar to the severing in a free pool case (see Figure S1). We show that both *L*_ss_ and the relaxation rate constant, which determines the time to steady state, are close to the expressions in the severing in a free pool case when 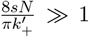 (see Supplementary Material for detailed derivations).

Next, we analyze the length trajectory obtained by stochastic simulations for both mechanisms. To compare the mechanisms, we chose parameters that result in the same steady-state filament length and found that the approach to reaching steady state and the length distribution at steady state are distinct (Figures 1E–1F) for the two cases. Filaments regulated by severing reached their steady-state length more quickly (Figure 1E) but exhibited a larger standard deviation and a skewed distribution in length (Figure 1F). At steady state, the scaling of vari-ance with the mean of the length distributions is distinct for the two mechanisms. In the case of constant disassembly in a limited pool, the variance scales inversely with the mean filament length (Figure 1G), while, in contrast, for cases where filaments can lose portions, the variance scales quadratically with the mean filament length (Figure 1H) [64].

As noted earlier, severing leads to larger fluctuations in filament lengths compared to constant disassembly. To study the differences, we analyzed the fluctuations in filament lengths at steady state (Figure 2A) using autocorrelation functions that measure the temporal correlations of the filament length fluctuations [65–67]. We observed that the autocorrelation function followed an exponentially decaying function, *e*^−*αt*^, for both mechanisms and noted that the decay in autocorrelation was faster for the severing mechanism than for constant disassembly (Figure 2B).

**Figure 2.**
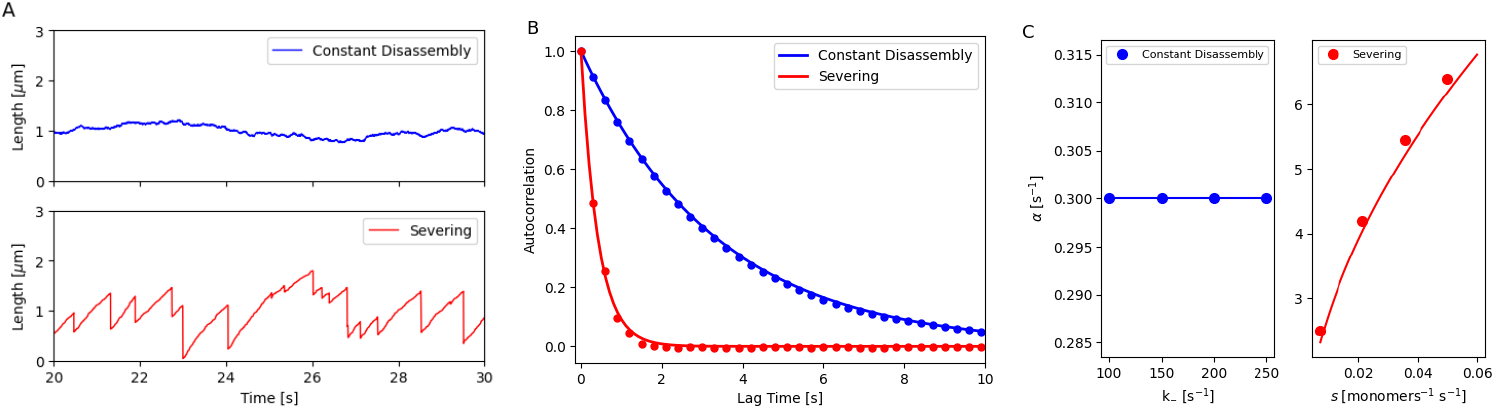
Analyzing fluctuations of a filament length assembled by (1) constant disassembly in a limited pool and (2) severing in a free pool. (A) Stochastic simulation of the steady-state length of a bare filament regulated by (1) constant disassembly in a limited pool (blue) and (2) severing in a free pool (red). (B) Autocorrelation of the steady-state filament length over time for the two mechanisms. Dots represent simulation results, while lines show analytical predictions (see Supplementary Material for detailed derivations). (C) Autocorrelation decay parameter *α* of steady-state filament length for different *k*_−_ values in constant disassembly and for *s* in severing mechanisms. Dots indicate simulation results, while lines represent analytical predictions (see Supplementary Material for detailed derivations).

Using the linear noise approximation, we compute a theoretical decay parameter *α* for each mechanism by linearizing the Langevin equation around the steady state and applying the Wiener-Khinchin theorem to derive the autocorrelation function [65–68] (see Supplementary Material). Remarkably, we find that, in the constant disassembly mechanism (using Equation 1), the decay parameter *α*_D_ is independent of *N* and *k*_−_, depending only on 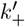. In contrast, for severing in a free pool mechanism, 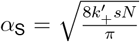, which depends on all three parameters (see Supplementary Material for detailed derivations for these two cases). Specifically, we find that the decay parameter has a distinct dependence on the disassembly parameters in the two models (Figure 2C).

The time for autocorrelation decay of filaments can be tuned by varying *α*_D_ or *α*_S_. In Figures S2 and 2B, we have chosen parameters that yield the same steady-state length, *L*_ss_ = 1 *µm*, for the two mechanisms with the same number of monomers *N*. For constant disassembly case, *L*_ss_ is a function of the ratio 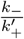, while for severing case, *L*_ss_ is a function of 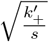. By choosing different values for 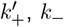 (constant disassembly), and *s* (with severing) to achieve a filament of 1 *µm* length, we observe that the autocorrelation decay can be made slower by decreasing *α*_D_ or *α*_S_.

In the Supplementary Material, we consider the growth of filaments (that can sever) in a limited pool and find that the decay parameter *α* is close to the one observed for severing (see Figure S1 for plots and Supplementary Material for detailed derivations).

### Assembly of Bundled Structures in a Pool

After modeling the growth dynamics of individual filaments, we extend our analysis to explore how the two disassembly mechanisms influence the growth dynamics of bundles. We assume that bundles consist of parallel individual filaments crosslinked together, with each fil-ament undergoing independent dynamics. As considered in a previous study [46], we model a bundle of *n* parallel filaments (with *n* = 6 in our study), where the length of the bundle, *L*, is defined as the maximum length of all filaments, i.e., *L* = max(*L*_*i*_) for *i* = 1, …, *n*.

In the main text, we consider two models: (1) *Constant disassembly in a free pool*, where filaments grow with an assembly rate 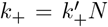 and disassemble at a constant rate *k*_−_; (2) *Severing in a free pool*, where filaments are subject to severing (see Figures 3A–3B). In Supplementary Material, we also consider growth in a limited pool (with and without severing) and provide a detailed comparison (see Figure S3).

**Figure 3.**
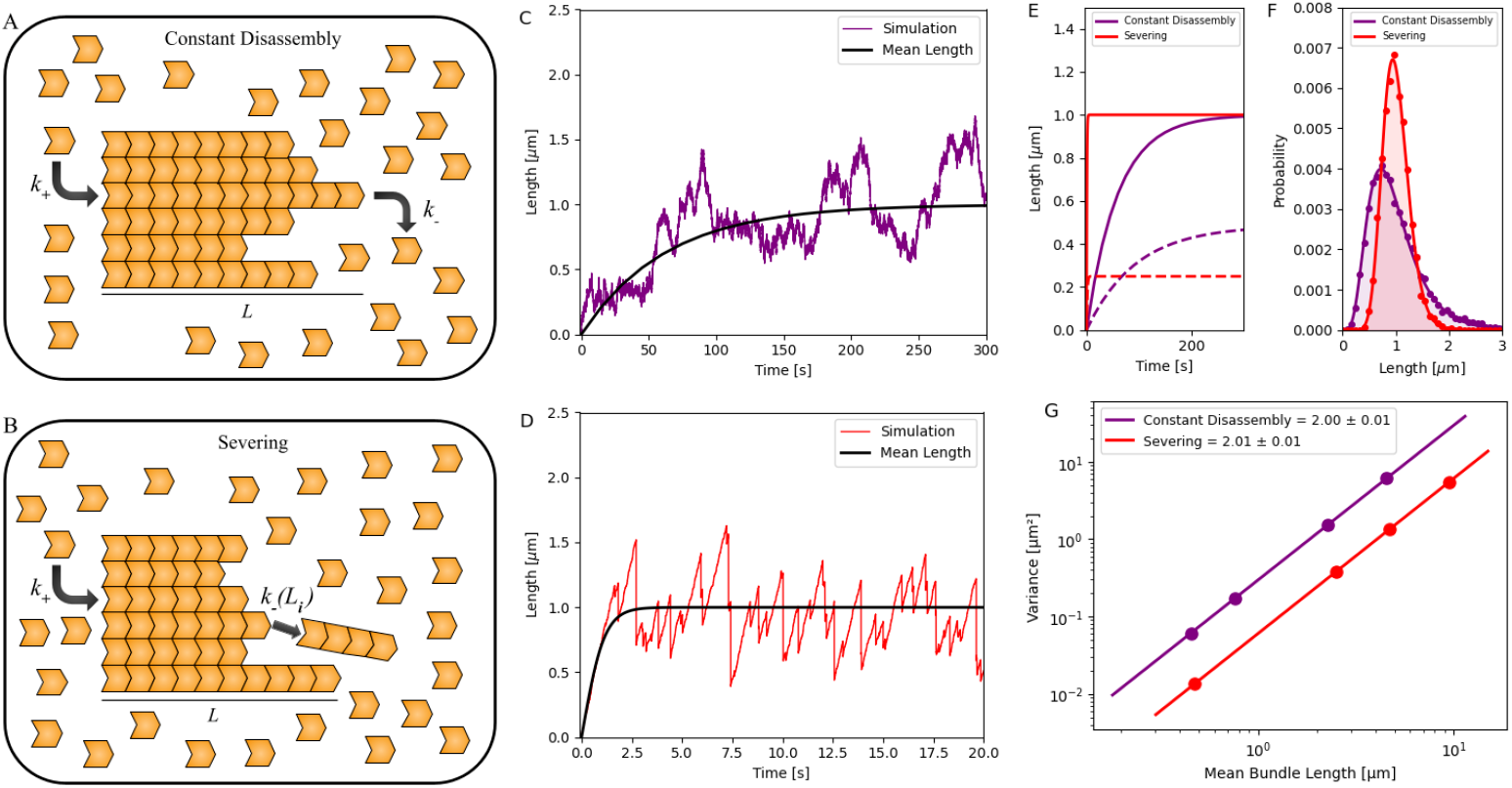
Growth dynamics of a bundle in a free pool with (1) constant disassembly and (2) severing. (A, B) Schematic showing the growth of a bundle in a free pool with (1) constant disassembly and (2) severing. (C, D) Stochastic simulations of bundle a bundle in a free pool with (1) constant disassembly (purple) and (2) severing (red). After an initial growth phase, the bundle length reaches a steady state. Simulation results are compared with fitted results, guided by an analytically derived calculation (black), in the Supplementary Material. (E) Stochastic simulations of the mean length dynamics (solid line) and standard deviation (dotted line) of bundle length, regulated by the two mechanisms, are plotted over time. (F) Probability distribution of the steady-state length of bundles regulated by the two mechanisms is shown, with dots representing simulation data and solid lines indicating analytical results [46] (see Supplementary Material). (G) Variance of filament length distributions is plotted against the mean filament lengths for the two cases, with variances presented on a log-log scale for both mechanisms. For panels (C–F), parameters used *N* = 100,000 monomers (each 4 nm in size), with 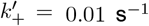; constant disassembly was modeled using *k*_−_ = 1009.7 s^−1^, and severing used *s* = 0.074 monomers^−1^ s^−1^. In panel (G), *N* = 200,000 monomers and 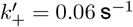 were used for all cases. For constant disassembly, *k*_−_ = {12025, 12050, 12150, 12250} s^−1^; for severing, *s* = {0.01, 0.04, 0.14, 4} monomers^−1^ s^−1^.

Using the Gillespie algorithm, we generate length trajectories and study the assembly dynamics of individual filaments within a bundle and the longest length at each instant (which we refer to as “bundle length”). We notice, only for certain combinations of parameters, that even though individual filament lengths may not reach a length distribution that is peaked at a specific size at steady state, the maximum filament length defining the bundle length does, as previously seen in [46] (see Figures 3C–3D). However, we also find other regimes where neither the filament lengths nor the bundle length reach a steady state. Using our simulations, we observe that in the constant disassembly mechanism, the bundle reaches a steady state only when the disassembly rate of individual filaments exceeds the assembly rate. In contrast, for assembly with severing, the bundle reaches a steady state across all regimes. We discuss these regimes further later in the manuscript.

Again, to compare different mechanisms, we chose parameters that would lead to a specific bundle size. We observed that while the two mechanisms can achieve a steady-state bundle length, the time scales associated with reaching it and their distributions at steady state are distinct (see Figures 3C–3F). We find that bundles regulated by the severing mechanism reach their steady-state length more quickly (Figure 3E) and with a lower standard deviation in length distribution compared to those assembled in a constant disassembly mechanism (Figure 3F). Note that this contrasts with the findings for a bare filament, where filaments regulated by sev-ering had a wider distribution at steady state (Figure 1F) than those in a constant disassembly.

Further, at steady state, both mechanisms predict that the variance of the distribution scales as the square of the mean length (Figure 3G), as reported in an earlier study [46], but with distinct intercepts. This suggests that while the models considered here can be candidate mechanisms for the assembly of bundles, the assembly kinetics might be a way to investigate whether processes like severing are involved in the assembly. We discuss this result later in the paper.

Next, we examined bundle length fluctuations and again used autocorrelation analysis of the steady-state bundle length (Figure 4A) to differentiate between the two mechanisms. The autocorrelation of bundle lengths followed an exponential decay, with the one assembled via severing having a faster decay than the Constant disassembly case (Figure 4B). We extracted an autocorrelation decay parameter *α* for both mechanisms and studied how it differs in its parameter dependence in each case (see Figure S4). For the severing mechanism, using linear expansion about the steady state, we find 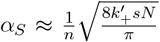, while we resort to simulations for extracting the parameter dependence in constant disassembly (Supplementary Material and Figure S4). Specifically, we find that *α* is sensitive to the assembly parameter 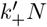 in constant disassembly, while increasing slowly in the case of severing (Figure 4C). Additionally, we find that in constant disassembly, the autocorrelation decay remains largely constant while the number of filaments is varied (*n* = 4 to 20), while it decays in the case of severing (Figure S4F).

**Figure 4.**
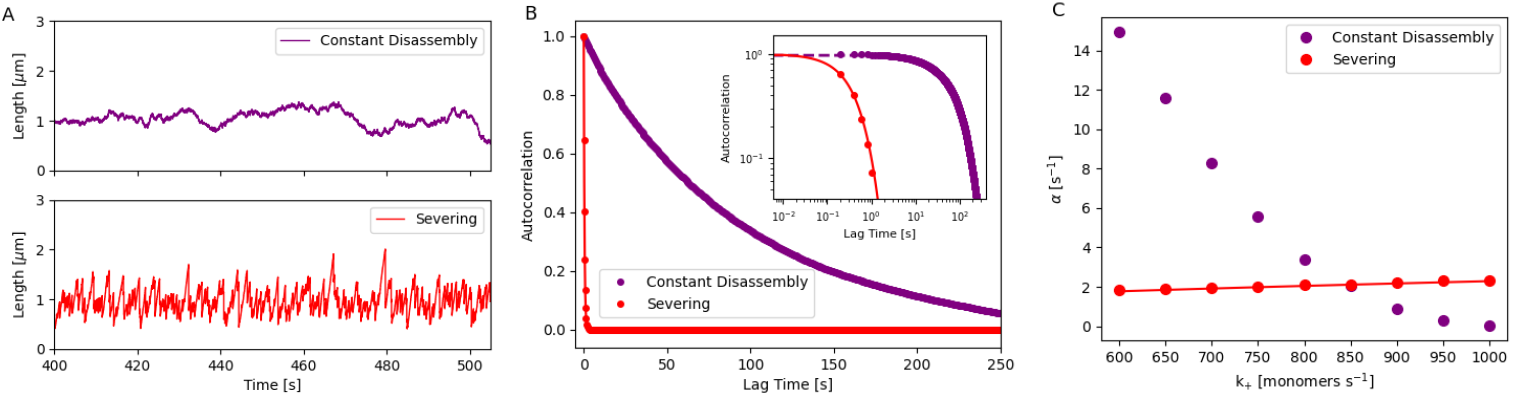
Analyzing fluctuations of bundle length assembled in a free pool with (1) constant disassembly and (2) severing. (A) Stochastic simulation of the steady-state bundle length assembled in a free pool with (1) constant disassembly (purple) and (2) severing (red). (B) Autocorrelation of steady-state bundle length over time for the two mechanisms. The inset figure shows the log-log plot. Dots represent simulation results, and solid lines show analytical predictions for the severing case (see Supplementary Material for detailed derivations). (C) Autocorrelation decay parameter *α* of the steady-state bundle length is analyzed for different *k*_+_ values in two mechanisms. Dots represent simulation results, while lines correspond to analytical predictions (refer to Supplementary Material for detailed derivations).

Other differences in parameter dependence of steady-state bundle length and the autocorelation decay parameter are discussed further in the Supplementary Figure S4. Additionally, we also assess the impact of different parameters on autocorrelation decay while maintaining a steady-state bundle length of *L*_ss_ = 1 *µ*m (Figure S2). In Supplementary Material, we also consider the effect of assembling a bundle in a limited pool and find that our results for autocorrelation decay *α* for the limited pool case are similar to the cases without (Figure S3).

### Co-assembly of Structures in a Shared Pool

Next, we examine the simultaneous assembly of filaments and bundles in a shared pool (Figures 5A–5B). We first consider *n* filaments (*n* = 1, 2, and 6 shown in Figure 5C) assembling under (1) constant disassembly in a limited pool, (2) severing in a free pool (Figure 5C), and (3) severing in a limited pool (Supplementary Material and Figure S5). As seen earlier, the assembly kinetics of the filaments are faster when severing is involved (see Figure S6). Additionally, we noticed interesting contrasts in their resulting steady-state distributions and fluctuations that can be used to discriminate between these mechanisms. We then analyze *b* bundles (*b* = 1, 2, and 6 shown in Figure 5D) assembling in a free pool under (1) constant disassembly and (2) severing (Figure 5D).

**Figure 5.**
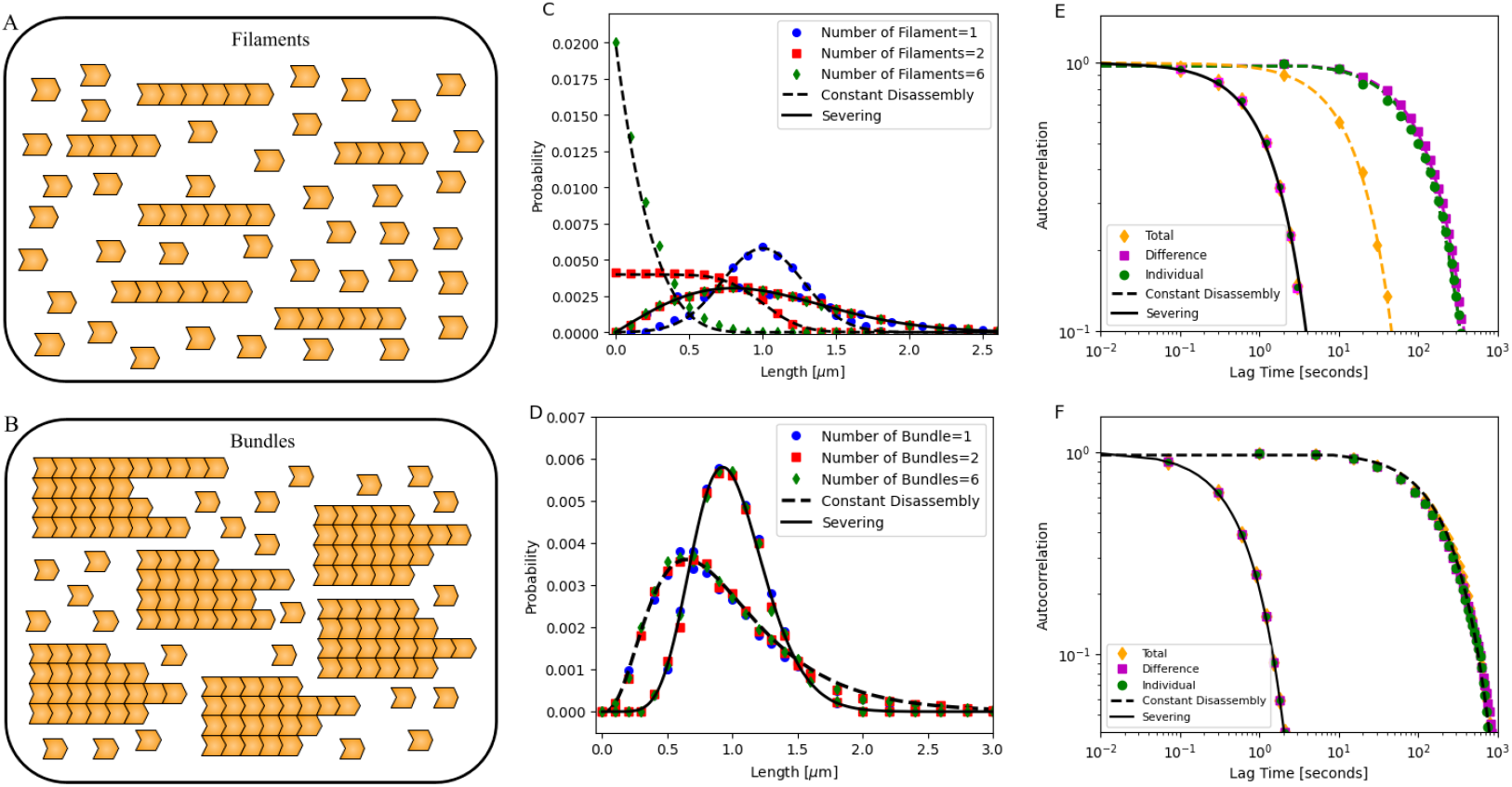
Probability distribution and autocorrelation of (1) multiple filaments regulated by constant disassembly in a limited pool and severing in a free pool, and (2) multiple bundles regulated by constant disassembly and severing in a free pool. (A–B) Schematic showing (A) *n* filaments assembling under constant disassembly in a limited pool and severing in a free pool, and (B) *b* bundles regulated by constant disassembly and severing in a free pool. (C) Steady-state length distributions for *n* filaments (*n* = 1, 2, 6) regulated by constant disassembly in a limited pool (dashed black lines) and by severing in a free pool (solid black lines). (D) Steady-state length distributions for *b* bundles (*b* = 1, 2, 6), each containing 4 filaments, regulated by constant disassembly in a free pool (dashed black lines) and by severing in a free pool (solid black lines). In both panels (C–D), the dots represent simulation results, while the lines indicate analytical predictions [32, 46, 61] (see Supplementary Material). (E) Autocorrelation functions for two co-assembling filaments under constant disassembly in a limited pool (dashed lines) and severing in a free pool (solid lines), showing individual filaments, their sum, and their difference. The plots are shown on a log-log scale, with dots representing simulation results, black solid and yellow dashed lines indicating analytical predictions, and other dashed lines corresponding to fitted data (see Supplementary Material). (F) Autocorrelation functions of steady-state lengths for two co-assembling bundles: individual bundles, their sum, and their difference, under both constant disassembly (dashed black lines) and severing in a free pool (solid black lines). Plots are shown on a log-log scale; dots represent simulation results, solid lines indicate analytical predictions, and dashed lines indicate fitted data (see Supplementary Material). Parameters: For filaments: *N* = 3000 (monomers, each 4 nm in size), 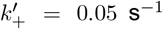, *k*_−_ = 137.5 s^−1^ (constant disassembly), *s* = 0.00375 monomer^−1^ s^−1^ (severing). For bundles with 4 filaments each, *N* = 100,000, 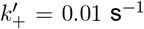, *k*_−_ = 1008.2 s^−1^ (constant disassembly), and *s* = 0.062 monomers^−1^ s^−1^ (severing).

As shown in our previous study [32], multiple filaments assembling under constant disassembly in a limited pool do not achieve a peaked steady-state distribution of lengths. While a single filament in a pool exhibits a steady-state length distribution centered around a mean, increasing the number of filaments in the shared pool broadens the distribution, with increasing variance. For six filaments, the distribution approaches an exponential form (Figure 5C). In contrast, in severing with free pool, co-assembling filaments attain a specific size, maintaining identical length distributions that peak at the same mean length with equal variance, regardless of filament number (Figure 5C). Interestingly, for severing in limited pool, we find that as the number of filaments *n* in a pool increases, both the mean and variance of the steady-state length distribution decrease, rather than peaking at the same mean with equal variance (see Supplementary Material for analytical derivations and Figure S5 for probability distribution plots). We also make predictions for how much of the pool is incorporated into each structure (see Figure S6). Interestingly, these specific predictions match with a recently published experimental study which examined the role of turnover mechanisms in the assembly of actin structures [69]. We discuss these results later. In comparison, bundles assembling in a shared pool with constant disassembly or severing exhibit identical and overlapping length distributions, irrespective of the number of bundles (*b* = 1, 2, and 6) co-assembling in the pool (Figure 5D), suggesting that these bundles are controlled independently. The assembly kinetics are once again faster with severing (see Supplementary Material and Figure S7).

Next, we analyze the fluctuations of two co-assembling filament and bundle lengths by computing their autocorrelation functions. In addition to the individual structure (filament or bundle) length autocorrelation, we show that the sum and difference of structure lengths can also reveal specific properties of each mechanism (see Supplementary Material for analytical derivations). For filaments assembling under constant disassembly in a limited pool, the sum of the two co-assembling filament lengths reaches a steady-state length and shows an exponentially decaying autocorrelation with a decay parameter 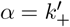 (see Supplementary Material for analytical derivations). In comparison, the individual filaments and their difference in lengths follow a random walk (as shown in [32]), with an autocorrelation function that decays much more slowly than the sum (Figure 5E). In contrast, with severing, filaments—including their sum and difference—exhibit identical and overlapping exponentially decaying autocorrelation functions with 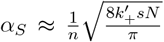 (Figure 5E and Supplementary Material for analytical derivations), suggesting that all filaments are assembling and controlled independently. Interestingly, in severing in a limited pool, the sum and difference of the lengths of the co-assembling filaments have distinct decay constants (see Supplementary Material for analytical derivations).

In the case of two bundles assembling in a free pool (with constant disassembly and severing), individual bundles, their sum, and difference exhibit identical and overlapping exponentially decaying autocorrelation functions, but each with distinct decay parameters (Figure 5F). As seen in the case of individual bundles, the decay parameter for severing is given by 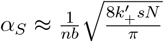 (see Supplementary Material). In contrast, simulations of the constant disassembly case show that the decay parameter is sensitive only to the assembly parameter 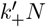. This suggests that with severing, co-assembling structures (filaments or bundles) are controlled independently. In Supplementary Material, we also consider the case of co-assembling bundles in a limited pool (with and without severing) and report minor differences in the decay parameter (Figure S5).

## DISCUSSION

In this study, we investigated the mechanisms that control the assembly of two types of structures— filaments and bundles—growing in a shared pool. We focused on two models of disassembly: one in which structures lose monomers at the ends at a constant rate, and another in which they lose parts of the filaments through severing, and we examined their effects on assembly dynamics and length fluctuations. We found that both models regulate structure size in distinct ways, and for a given length, assembly dynamics accelerate when severing is involved. While severing results in a wider length distribution at steady state for filaments, bundles exhibit a distribution with lower variance. Additionally, we found that severing can regulate multiple filaments and bundles assembling simultaneously in a shared pool, constraining them to a specific size. Furthermore, by analyzing length fluctuations at steady state, we observed that, in general, the decay in the autocorrelation function is faster when severing is included. Significantly, our study identifies parameters that can be tuned in experiments to modify assembly kinetics, control the decay of autocorrelations, and help distinguish between different disassembly mechanisms. In this section, we discuss these results.

### Severing Can Accelerate and Modulate Relaxation Dynamics

Across all mechanisms considered to create a structure of a given length, we find that incorporating severing accelerates the assembly of both types of structures. We also identify key parameters that can be tuned to enhance their assembly from a shared monomer pool. As described in the Supplementary Material, for a single filament undergoing constant disassembly, the growth parameter 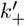 solely determines the relaxation rate to steady state. In contrast, for bundles assembling from a shared pool, our simulations show that the relaxation rate is sensitive to the product 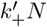 (Figure S4G). However, when severing is included, the relaxation dynamics of both bare filaments and bundles are governed by a combination of parameters, 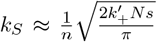 (Supplementary Material). Interestingly, with severing, the relaxation rate in-creases only slightly with 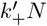, in contrast to the monotonic decay observed with constant disassembly (see Supplementary Material and Figure S4G). These distinct dependencies offer a stringent test for distinguishing whether the assembly of structures involves large disassembly events such as severing.

### Disassembly Mechanisms Can Cause Differential Regulation of Co-assembling Structures in a Shared Pool

We found that severing regulates the lengths of co-assembling filaments and bundles in a shared monomer pool. Our analysis also predicts how assembly kinetics and total monomer incorporation vary with the number of co-assembling structures. In general, we observe that (1) severing accelerates assembly, and (2) increasing the number of structures slows down assembly (Figures S6 and S7).

For filaments assembling in a limited pool, individual filament lengths at steady state decrease as the number of filaments increases (Figure S6A). When severing is included, size variability is reduced, and filament lengths converge, regardless of their number, in the limit of a free monomer pool (Figures S6B–C). Moreover, under constant disassembly, the total monomer content in all filaments remains independent of filament number (Figure S6D), whereas with severing, total monomer incorporation scales with the number of filaments (Figures S6E–F). Interestingly, these predictions align with experimental observations from recent work on actin “comet tails” assembled in microwells with limited monomer supply [69]. In this study, comet tails were reported to reach a characteristic size even with minimal turnover, with a slow growth. Upon accelerating turnover by the addition of ADF/cofilin and capping proteins, growth rates increased significantly. Furthermore, total polymerized actin remained constant across varying numbers of comets without turnover but increased with the number when turnover was present, similar to what we predict.

### Fluctuation Analysis Can Be Used to Distinguish Mechanisms of Assembly

Although all assembly mechanisms considered here eventually reach steady state, the length fluctuations around this state differ. In particular, we find that the decay of autocorrelation functions is consistently faster when severing is involved. For both bare filaments and bundles regulated by severing, we derived that the decay parameter is given by 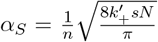. In contrast, for bare filaments regulated by constant disassembly within a limited pool, *α* is solely dependent on 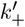, while for bundles, *α* is highly sensitive to the assembly parameter 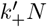. Extracting this decay parameter from the fluctuations at steady state, and studying its dependence on different parameters like 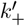 or *n*, can thus be used to contrast severing-based mechanisms with other length-control mechanisms. These insights provide a framework for designing experiments aimed at investigating the precise role that mechanisms like severing play in the assembly of cytoskeletal structures. This approach can be extended to other size-control mechanisms, and examining autocorrelation as a function of model-specific parameters could thus offer a valuable method for distinguishing between these mechanisms.

### Assembly Dynamics Can Be Used to Investigate Length-Control Mechanisms in Bundles

A recent study reported that the scaling of variance with the mean in many bundled structures follows a quadratic relationship [46]. This study also proposed an elegant mathematical model showing that if each filament length in a bundle is assumed to be exponentially distributed, then the bundle length (defined as the longest filament within the bundle) is controlled. The resulting distribution exhibits a quadratic relationship between variance and mean. Our study, which simulates filaments in a shared pool using assembly and disassembly rates without assuming an exponential length distribution for filaments in the bundle, confirmed this finding: at steady state, all mechanisms of bundle assembly considered here exhibited a quadratic scaling relationship between variance and mean. Interestingly, we find that bundles regulated by severing have a different intercept in the variance vs. mean relationship, as seen in Figure 3G.

Further, using our simulation scheme, we identified distinct parameter regimes where bundles achieved a steady state. For constant disassembly in a free pool, steady-state bundle lengths were achieved only when the disassembly rate exceeded the assembly rate. However, with severing, this condition was not necessary, and the bundles reached a steady state under all conditions. Additionally, we found that assembly kinetics were faster when severing was involved. These findings suggest that a class of models could explain the variance scaling observed in experimental measurements of bundles, and scrutinizing the parameter dependence of assembly kinetics in each case may provide insights into the role of accessory mechanisms, such as severing, in bundle assembly.

To compare the effects of different disassembly mechanisms, we examined two limiting cases: monomer dissociation occurring one unit at a time, and severing with large dissociation events. We assumed that severing occurs uniformly along the filament length. This assumption is supported by in vitro studies on microtubules, where the severing protein Spastin has been shown to sever uniformly along the lattice [70]. In contrast, actin severing proteins such as cofilin exhibit binding preferences that depend on the nucleotide state of actin monomers which may modulate the disassembly rates between two limits considered here. Furthermore, efficient and rapid actin severing is often mediated by a combination of proteins—including cofilin, coronin, and Aip1—which act cooperatively to enhance disassembly and turnover [71]. We further assume that the severed filaments are capped and eventually disassemble to become part of the monomer pool.

Despite these biological complexities, our qualitative conclusion—that severing leads to faster relaxation rates and autocorrelation decay—holds for any mechanism involving large dissociation events. The modeling framework presented here can be extended to incorporate more detailed biophysical features of disassembly processes, allowing for a deeper understanding of how such mechanisms influence the growth kinetics and fluctuations of filamentous structures.

In cells, multiple regulatory mechanisms likely act in concert to control organelle and cytoskeletal structure assembly. A systematic understanding of how these mechanisms interact is crucial to decoding the design principles of assembly. In this study, we analysed two key disassembly pathways and their effects on the assembly of bare filaments and bundles in a shared resource pool. Our findings highlight specific parameters that can be tuned to modulate assembly kinetics, structure sizes, and fluctuations. Ultimately, such approaches could reveal how cells achieve precise and robust control over their internal organization.

## Acknowledgements

We extend our gratitude to the members of the RIT Soft Matter Group—Peter Miller, Nishant Malik, and Moumita Das and her lab—for their valuable and stimulating discussions. This research was supported by the National Institute of General Medical Sciences (NIGMS) of the National Institutes of Health (NIH) under Award Number R35GM147556 (M.S.A.M., M.C., L.M.).

## Supplementary Material

### Derivation of Mean Length Dynamics for Bare Filament and Bundled Structures

#### Bare Filament

##### Assembly of a bare filament with severing in a limited pool

Solving Equation 2 in the main text, with a growth rate 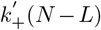, a severing rate *s*, and initial conditions *t* = 0, *L* = 0, the time evolution of filament length is given by *L*(*t*) = *L*_ss_ tanh(*k*_LPS_*t*), where the steady-state length and relaxation rate are given by 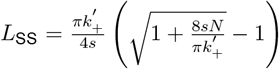 and 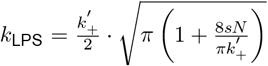, respectively. This equation is plotted in Figure S1B.

For large values of *N*, where 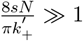, the relaxation rate simplifies to *k*_LPS_ ≈ *k*_S_, which is equivalent to the rate *k*_S_ found in the severing scenario with a free monomer pool.

#### Bundled Structure

##### Assembly of a bundle with severing in a free pool

Here, the length dynamics of an individual filament in a bundle of *n* filaments are described by

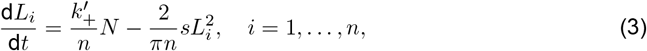

where 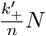 and 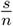 represent the Identical growth and severing rates for individual filaments, while 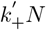 and *s* represent the growth and severing rates for the sum total of *n* filaments.

Assuming equal filament lengths *L*_*i*_ = *L*_avg_, the average length evolves as

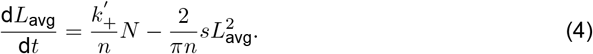

With initial conditions at *t* = 0, *L*_avg_ = 0, the time evolution of filament length is given by*L*_avg_(*t*) = *L*_ss_ tanh(*k*_S_*t*), where the steady-state length and relaxation rate are 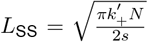 and 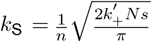, respectively. This equation is used to calculate the autocorrelation function of aver-age filament length at steady state.

##### Assembly of a bundle with severing in a limited pool

Here, the length dynamics of an individual filament in a bundle of *n* filaments are described by

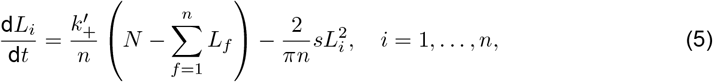

where, 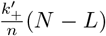 and 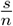 denote the identical growth and severing rates of the individual fila-ments, whereas 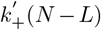 and *s* correspond to the growth and severing rates for the sum total of *n* filaments. Assuming equal filament lengths *L*_*i*_ = *L*_avg_, the average length evolves as

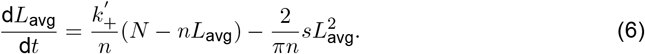

Starting with the initial condition *L*_avg_(0) = 0, the filament length evolves over time according to *L*_avg_(*t*) = *L*_SS_ tanh(*k*_LPS_*t*), where the steady-state length and the relaxation rate are given by 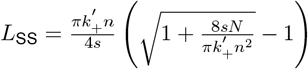 and 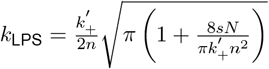, respectively. This expression is used to calculate the autocorrelation function of the average filament length at steady-state.

For large *N* such that 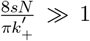, the relaxation rate *k*_LPS_ approximates *k*_S_, matching the relaxation rate observed in the severing scenario with a free monomer pool.

### Probability Distribution of Lengths of Bare Filaments and Bundled Structures

The detailed analytical derivations for the probability distributions of bare filaments and the coassembly of multiple filaments under the constant disassembly mechanism within a limited pool are provided in our previous work [32]. The derivations corresponding to severing dynamics in a free pool are presented in our earlier studies [61, 64]. Furthermore, the analytical derivations for the probability distributions of bundles undergoing constant disassembly and severing in a free pool is discussed in [46]. In the present work, we extend these analyses by deriving the length probability distributions for both bare filaments and bundles regulated by severing within a limited pool.

### Probability Distribution of Lengths of Bare Filaments Under Constant Disassembly in a Limited Pool

The master equation governing the probability *P* (*L, t*) for a filament of length *L* under a limited pool with severing is:

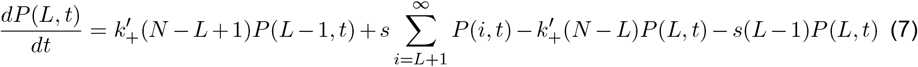

At steady state, where 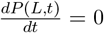, and applying the normalization condition 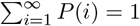, we define 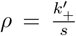. Using the recursive method from our previous work [61], the probability distribution for *L* ≥ 0 is given by

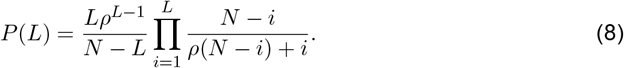

This equation is plotted in Figure S1D. To simplify, we rewrite the product and approximate it for large *ρ*(*N* − *i*). For *L* ≪ *N*, this yields the approximate probability distribution 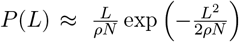. Substituting 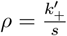, we obtain the final form:

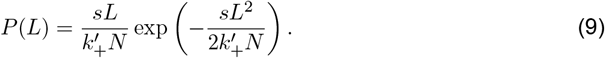

### Probability Distribution of Individual Filament Lengths in a Bundle of *n* Filaments Regulated by Severing in a Limited Pool

First, we consider two filaments, *L*_1_ and *L*_2_, growing in the pool. The master equations for these two filaments are given by:

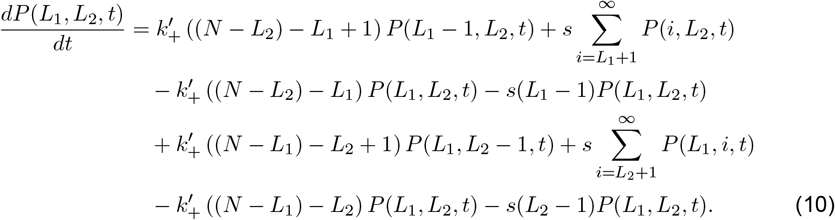

When the total number of monomers *N* is very large, the influence of one filament’s average length on the average length of other filaments becomes negligible. As a result, at any given time, the length of one filament (say *L*_1_) depends primarily on the available number of free monomers, which is given by *N* ^∗^ − *L*_1_ = *N* − ⟨*L*_2_⟩ − *L*_1_, where *N* ^∗^ = *N* − ⟨*L*_2_⟩ and ⟨*L*_2_⟩ is the steady-state average length of *L*_2_. Under these conditions, the master equation for individual filament *L*_1_ simplifies to:

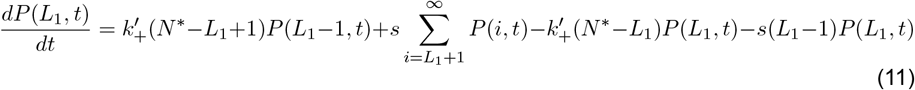

At steady state, i.e., when 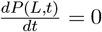, and using the normalization condition 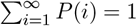, we define 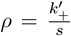. Following the recursive approach outlined in our previous work [61], The resulting probability distribution for *L*_1_ is then:

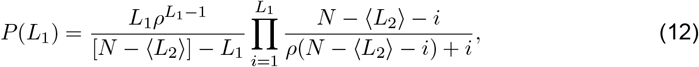

This equation can be further simplified in a manner similar to Equation 9 as follows:

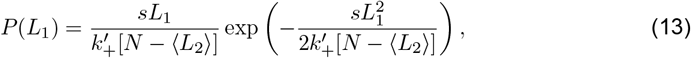

To find the average filament length ⟨*L*_1_⟩, we use the expectation formula 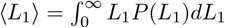. Substituting *P* (*L*_1_) and simplifying, the integral reduces to a Gaussian form. Using the known integral 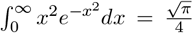, we get 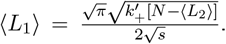. Assuming ⟨*L* ⟩ = ⟨*L* ⟩ = ⟨*L*⟩, the steady-state length satisfies 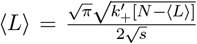. Solving this quadratic equation for the positive root, we obtain

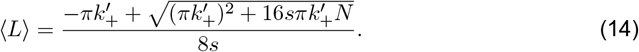

Similarly, when there are *n* filaments *L*_1_, *L*_2_, …, *L*_*n*_ present in the pool and regulated by severing in a limited pool mechanism, we can generalize Equation 12 as:

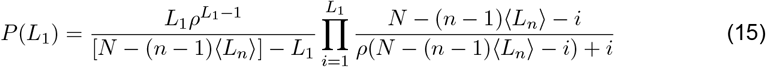

where,

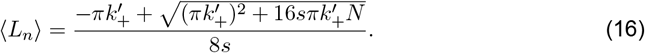

This equation is plotted in Figure S5A.

### Probability Distribution of the Bundle Length Consisting of *n* Filaments Regulated by Severing in a Limited Pool

For a bundle consisting of *n* filaments, each of length *l*_*i*_, the filament lengths follow the distribution

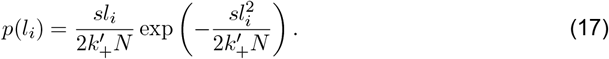

The cumulative probability that a filament length is less than *L* is 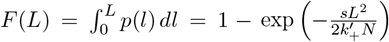, and therefore, the probability that all *n* filaments have lengths less than *L* is 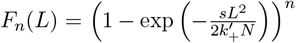. The probability density function for the longest filament length is given by the derivative of *F*_*n*_(*L*),

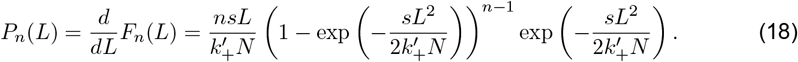

This equation is plotted in Figure S3F.

### Autocorrelation Decay Parameter for Bare Filaments, Bundled Structures, and Co-assembly of Structures

In order to calculate the autocorrelation decay, we use Langevin equations associated with different growth mechanisms. For a univariate stochastic process *y*_*t*_, the autocorrelation at lag *k* is given by 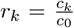, where 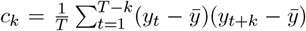 is the lag-*k* covariance, and *c*_0_ is the sample variance of the time series [67]. According to the Wiener–Khinchin theorem, the autocorrelation function of a stationary process is also the inverse Fourier transform of its power spectral density [65, 66, 68]. Using the techniques outlined in [65, 66, 68], we derive the expressions for the autocorrelation function corresponding to the various growth mechanisms considered here.

#### Bare Filament

##### Assembly of a bare filament with constant disassembly in a limited pool

Here, the dynamics of filament length are governed by the following stochastic differential equation:

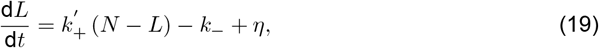

where *L* denotes the filament length, and *η* represents a stochastic noise term associated with its dynamics. At steady state, setting the time derivative and the average noise to zero yields the total filament length 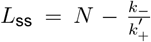. To study fluctuations around the steady state, we introduce a small perturbation *L* = *L*_ss_ + Δ*L*, and substitute this into the dynamic equation for filament length. Since 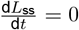, and using the expression for *L*_ss_, we obtain:

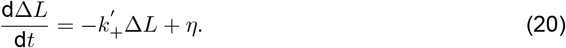

Next, taking the Fourier transform of the dynamical equation yields 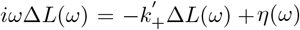, where Δ*L*(*ω*) and *η*(*ω*) are the Fourier transforms of the length fluctuation and the noise, respectively. Rearranging, we obtain 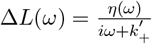, leading to a power spectrum of 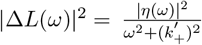. Assuming delta-correlated noise, 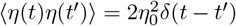, which implies a flat noise spectrum 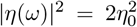, we find that the power spectrum of length fluctuations is given by 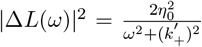. By the Wiener-Khinchin theorem, the autocorrelation function is the in-verse Fourier transform of the power spectrum, yielding 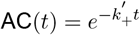, with an autocorrelation decay parameter 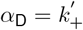. This equation is plotted in Figures 2B–2C and Figure S2A.

The same technique has been employed to determine the autocorrelation decay for the other growth mechanisms as well.

##### Assembly of a bare filament with severing in a free pool

The filament length dynamics are described by the stochastic differential equation

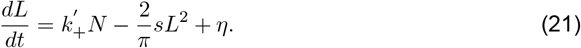

Using the methodology outlined above, the autocorrelation decay rate is given by 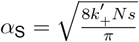.The corresponding plot of this equation is shown in Figures 2B–2C and Figure S2B.

##### Assembly of a bare filament with severing in a limited pool

The filament length dynamics are described by the stochastic differential equation

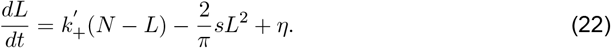

Applying the previously described methodology, we obtain the autocorrelation decay rate as 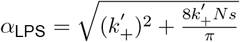. This equation is plotted in Figure S1F.

#### Bundled Structure

##### Assembly of a bundle with severing in a free pool

We analyze the dynamics of a bundle consisting of *n* filaments, where the length evolution of an individual filament is governed by

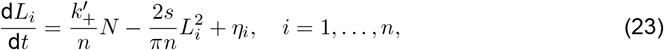

with 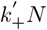 representing the polymerization rate and *s* denoting the severing rate of the total filament length. Assuming all filaments maintain equal lengths, i.e., *L*_*i*_ = *L*_avg_, the dynamics reduce to the following equation for the average filament length:

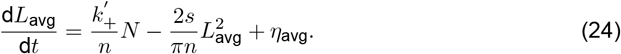

Using the same technique described above, we obtain the autocorrelation decay parameter for the average filament length as 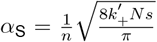. This equation for the average filament length closely agrees with the simulated autocorrelation decay parameter of the bundle length, and the corresponding plots are shown in Figures 4B–4C, S2D, and S4E–S4F.

##### Assembly of a bundle with severing in a limited pool

Here the length evolution of an individual filament is governed by

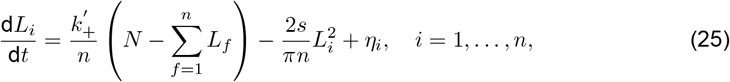

with 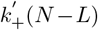 representing the polymerization rate and *s* denoting the severing rate of the total filament length. Assuming all filaments maintain equal lengths, i.e., *L*_*i*_ = *L*_avg_, the dynamics reduce to the following equation for the average filament length:

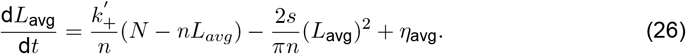

Using the same technique described above, we obtain the autocorrelation decay parameter for the average filament length as 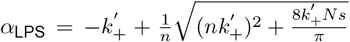. This equation for the average filament length closely agrees with the simulated autocorrelation decay parameter of the bundle length, and is plotted in Figure S3H.

#### Co-assembly of Structures

##### Assembly of multiple filaments regulated by constant disassembly in a limited pool

We consider *n* filaments, where the total growth rate of all filaments is given by 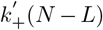, and the disassembly rate is given by *k*_−_. The dynamics of the filament lengths are described by:

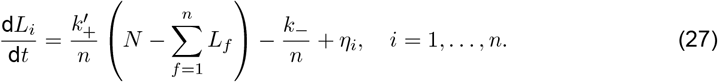

Assuming all filaments in the bundle have equal length, denoted by *L*_1_ = *L*_2_ = · · · = *L*_*n*_ = *L*_avg_, where *L*_avg_ represents the average length of the *n* filaments, the total filament length 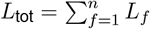 and the total noise term 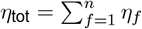; the time evolution of the total filament length is then described by:

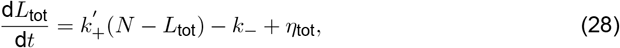

By applying the previously outlined method, the autocorrelation decay parameter for the total filament length is found to be 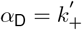. This equation is plotted in Figure 5E.

##### Assembly of multiple filaments regulated by severing in a free pool

We consider a system of *n* filaments, where the length of each filament evolves according to

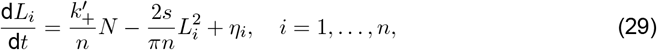

with 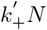 representing the total polymerization rate and *s* the severing rate. Assuming all fila-ments remain of equal length, *L*_*i*_ = *L*_avg_, the dynamics of the total filament length 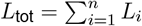 simplify to

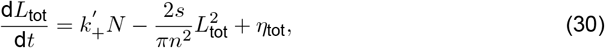

where 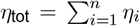 represents the cumulative noise from all filaments. Applying the same Fourier-based analysis as described earlier, we obtain the autocorrelation decay rate for the total filament length as 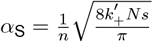. This equation is plotted in Figure 5E.

To analyze the relative dynamics between filaments, we consider the length difference between any two filaments *i* and *j*, denoted by *L*_diff_ = *L*_*ij*_ = *L*_*i*_ − *L*_*j*_. By subtracting the corresponding equations for *L*_*i*_ and *L*_*j*_, the dynamics of the filament length difference are given by:

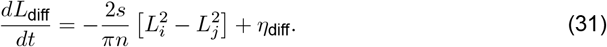

Applying the same method described above, we derive the autocorrelation decay parameter for the filament length difference, which is the same as the expression derived for the total filament length: 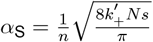. This equation is plotted in Figure 5E.

##### Assembly of multiple filaments regulated by severing in a limited pool

Here, we analyze a system consisting of *n* filaments, where the growth of each filament is regulated by both a shared monomer pool and severing. The evolution equation for the length of the *i*^th^ filament is given by:

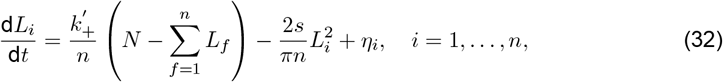

Where 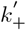 is the polymerization rate per filament and *s* denotes the severing rate. Assuming uniform filament lengths, i.e., *L*_*i*_ = *L*_avg_ for all *i*, the total filament length 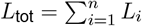 follows the reduced form:

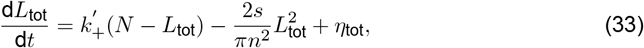

with 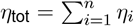 capturing the collective noise across all filaments. By applying the Fourier-based approach used in previous sections, the autocorrelation decay rate for the total filament length is found to be 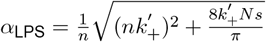.

To further understand the internal dynamics of the system, we consider the difference in length between two filaments, defined as *L*_diff_ = *L*_*i*_ − *L*_*j*_. Subtracting their individual length evolution equations yields the following expression:

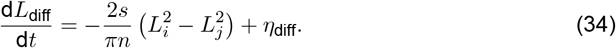

Applying the same analytical technique, we derive the autocorrelation decay rate for this length difference as 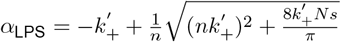.

##### Assembly of multiple bundles regulated by severing

We consider *b* bundles with *n* filaments each, resulting in a total of *nb* filaments. Using the Fourier-based technique outlined above, the autocorrelation decay parameter for a free pool is given by 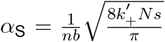. This equation is plotted in Figure 5F. For a limited pool, the decay parameter becomes 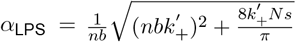, which is shown in Figure S5B.

#### Simulation Protocol

We employed Gillespie stochastic simulations to model the growth of bare filaments and bundles. A bundle is defined as a collection of linear, parallel filaments, with its length determined by the maximum length among them. For both bare filaments and bundles, simulations begin with all filaments initialized at zero length. At each simulation step, a possible transition— corresponding to a change in filament length—is randomly selected with a probability proportional to its associated rate. The time interval between transitions is drawn from an exponential distribution, whose rate parameter equals the sum of all possible transition rates from the current state. This process is repeated until the system reaches a steady-state length distribution. To ensure statistical reliability, we generate multiple independent stochastic trajectories and compute the steady-state distributions of filament and bundle lengths by averaging over these realizations.

To analyze fluctuations in the steady-state time series of filament and bundle lengths, we computed the autocorrelation using MATLAB’s autocorr function.

## Supplemental Results

**Figure S1.**
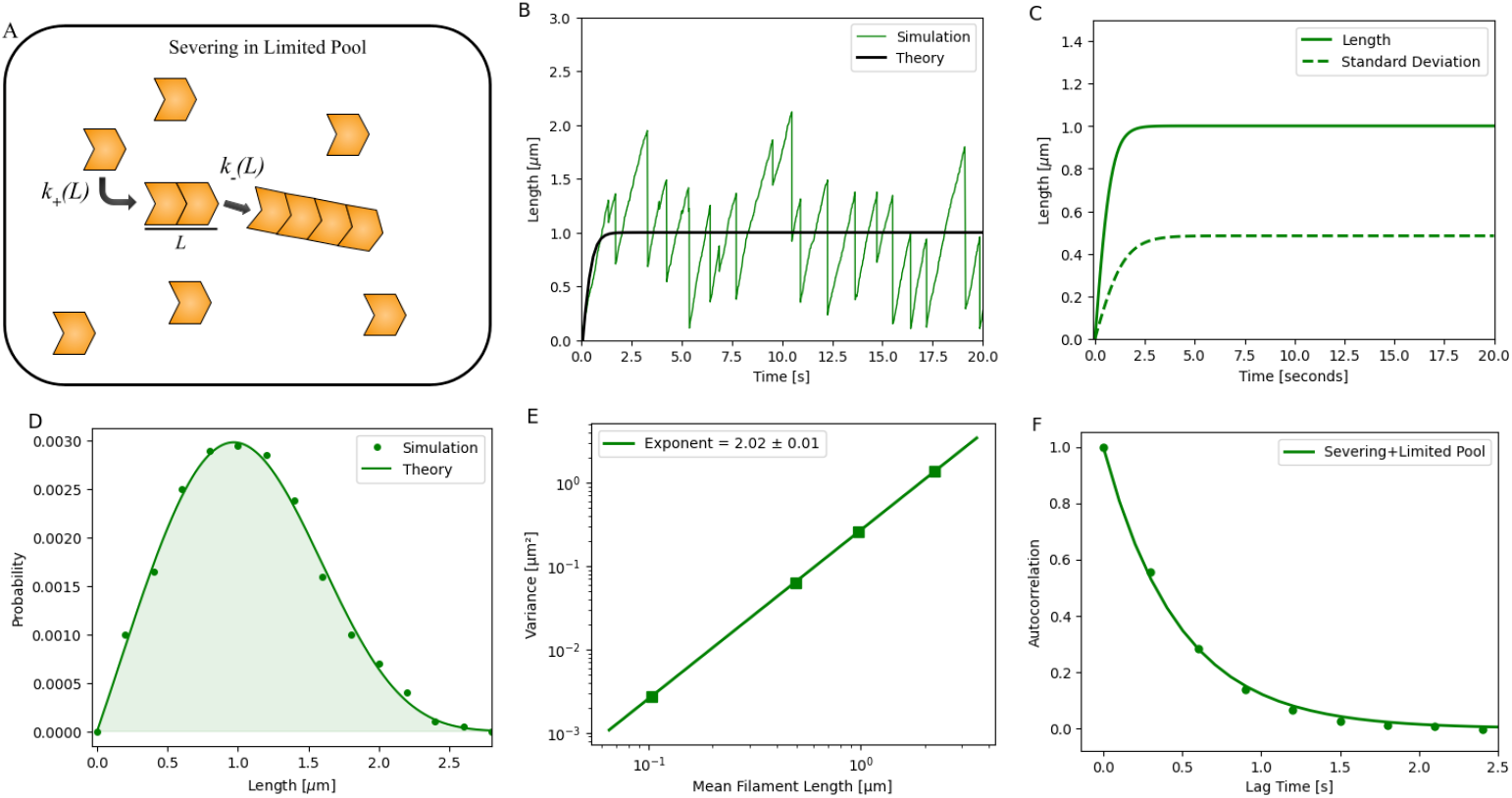
Growth dynamics of a bare filament under severing in a limited pool. (A) Schematic illustrating the growth of a bare filament. (B) Stochastic simulations of filament growth trajectories. Simulation results are overlaid with analytical predictions (black), as described in the Supplementary Material. (C) Mean length (solid lines) and standard deviation (dashed lines) of bare filaments, calculated from simulations, plotted as a function of time. (D) Probability distribution of steady-state filament lengths. Dots represent simulation data; solid lines indicate analytical predictions (see Supplementary Material). (E) Variance of filament length distributions plotted against the corresponding mean filament lengths (log-log scale). (F) Autocorrelation of steady-state filament length over time. Dots represent simulation results; lines show analytical predictions (see Supplementary Material for detailed derivations). Parameters: *N* = 1000 monomers (each 4 nm in size), 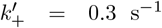, and *s* = 0.00565 monomers^−1^ s^−1^. For panel (E), different values of 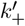 were selected to obtain distinct mean filament lengths, with *N* = 10000: 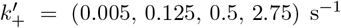, 0.125, 0.5, 2.75) s^−1^ and *s* = 0.125 monomers^−1^ s^−1^.

**Figure S2.**
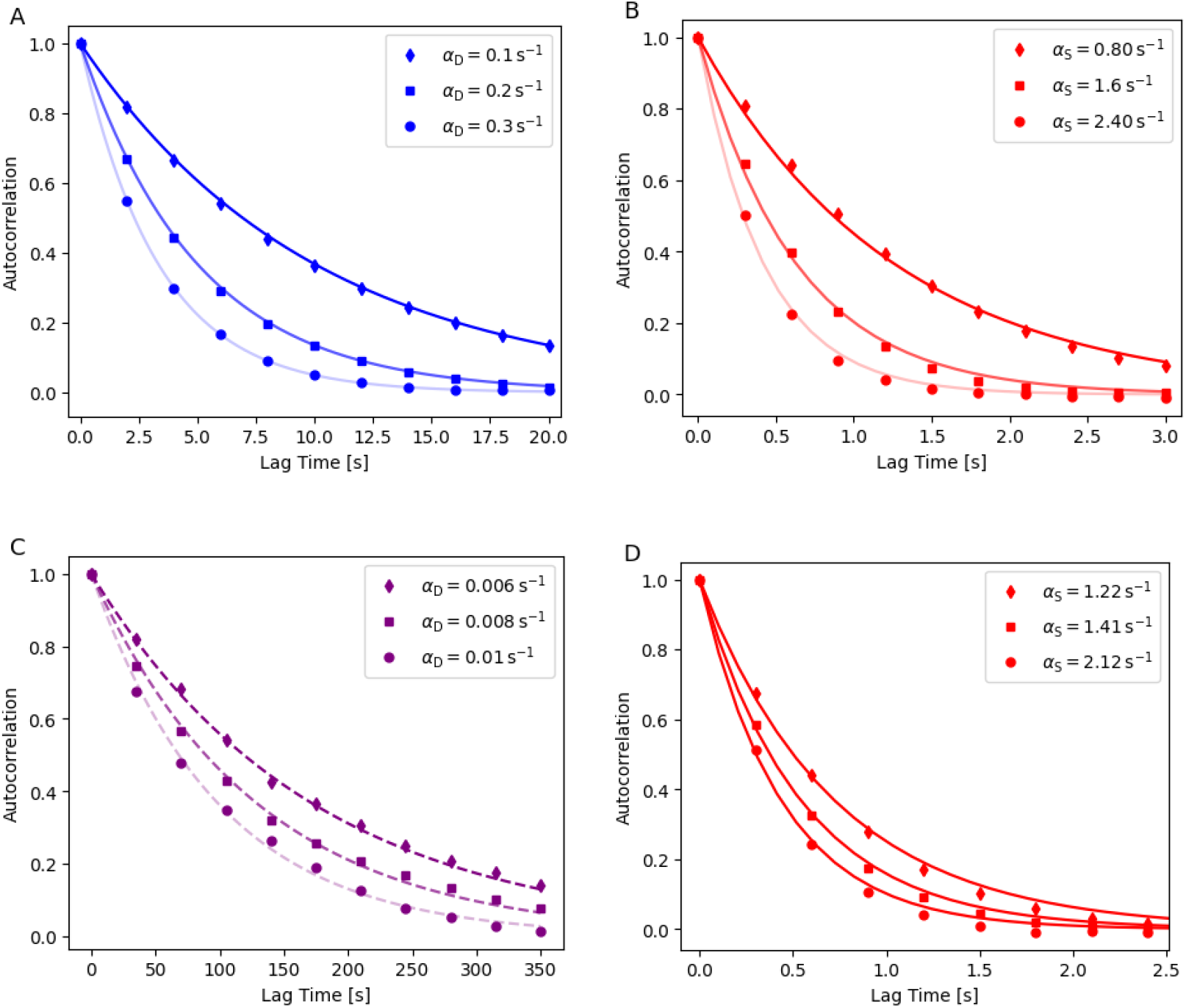
Autocorrelation analysis of steady-state length fluctuations in single filaments and bundles under various assembly-disassembly mechanisms. (A–B) Filaments: Autocorrelation of filament length over time for different values of *α*, with *N* = 1000 and mean filament length maintained near 1 *µ*m using (A) constant disassembly in a limited pool and (B) severing in a free pool. (C–D) Bundles: Corresponding autocorrelation plots for bundle length with *N* = 100000, again maintaining ∼1 *µ*m mean filament length, under (C) constant disassembly in a free pool and (D) severing in a free pool. Dots represent simulation data; solid lines are analytical predictions, and the dashed line is the fitted result (see Supplementary Material for derivation details).

**Figure S3.**
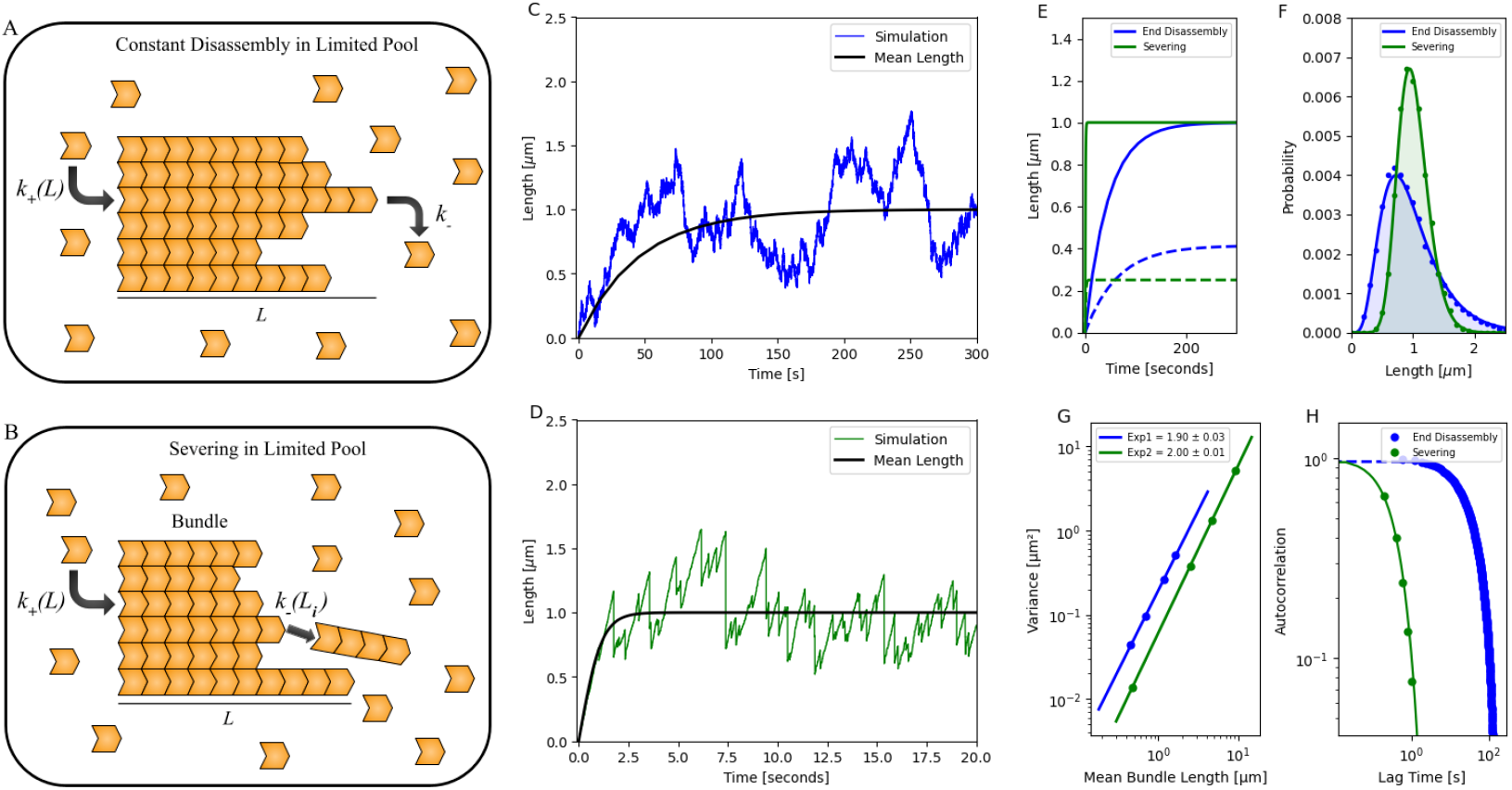
Growth dynamics of a bundle in a limited pool with (1) constant disassembly and (2) severing. (A, B) Schematics showing the growth of a bundle in a limited pool with: (1) constant disassembly and (2) severing. (C, D) Stochastic simulations of bundle growth regulated limited pool with (1) constant disassembly and (2) severing. After an initial growth phase, bundle length reaches a steady state. Simulation results are compared with fitted results (black), guided by analytical calculations described in the Supplementary Material. (E) Stochastic simulations of the mean bundle length (solid line) and standard deviation (dotted line) over time for both mechanisms. (F) Probability distribution of steady-state bundle lengths for the two mechanisms. Dots indicate simulation data, and solid lines represent analytical results (see Supplementary Material). (G) Variance of bundle length distributions plotted against the mean bundle lengths. Both mechanisms are shown on a log-log scale. (H) Autocorrelation of steady-state bundle length over time for the two mechanisms, shown on a log-log scale. Dots represent simulation results, solid lines indicate analytical predictions, and dashed lines denote fitted data (see Supplementary Material). Parameters for panels (C–F): *N* = 100,000 monomers (each 4 nm in size), with 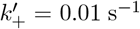. For the limited pool with end disassembly, *k*_−_ = 1002.8 s^−1^; for the limited pool with severing, *s* = 0.074 monomers^−1^ s^−1^. In panel (G), simulations were performed with *N* = 200,000 monomers and a constant assembly rate 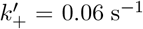 across all cases. For the limited pool with end disassembly, disassembly rates were *k*_−_ = {12005, 12050, 12150, 12250} s^−1^; for the limited pool with severing, severing rates were *s* = {0.01, 0.04, 0.14, 4} monomers^−1^ s^−1^.

**Figure S4.**
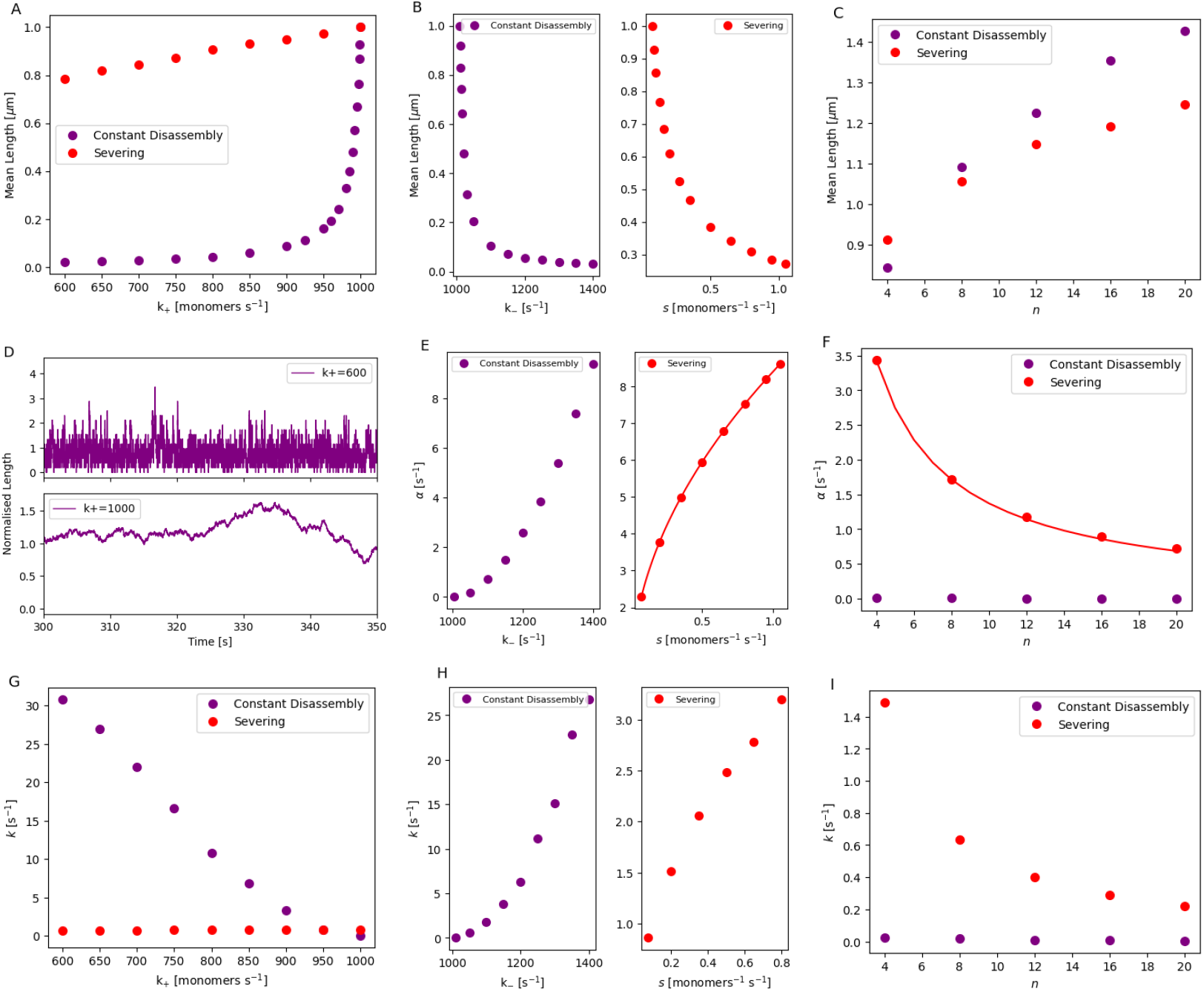
Parameter dependence of length, autocorrelation decay, and relaxation rate for bundles in a free monomer pool regulated by (1) constant disassembly and (2) severing. (A–C) Steady-state bundle length for the two mechanisms plotted as a function of *k*_+_, *k*_−_, and *n*, respectively, with other parameters held constant. (D) Simulated time trajectory of the steady-state bundle length regulated by simple assembly for *k*_+_ = 600 and 1000, respectively, with other parameters held constant. (E–F) Autocorrelation decay parameter of the steady-state bundle length for the two mechanisms plotted as a function of *k*_−_, and *n*, respectively. Dots represent simulation results, and lines indicate analytical predictions (see Supplementary Material for detailed derivations). (G–I) Relaxation rate toward the steady-state bundle length for the two mechanisms plotted as a function of *k*_+_, *k*_−_, and *n*, respectively, with other parameters held constant. Parameters: *N* = 100,000 monomers (each 4 nm in size), 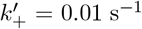; for simple assembly, *k*_−_ = 1010 s^−1^; for the free pool with severing, *s* = 0.074 monomers^−1^ s^−1^.

**Figure S5.**
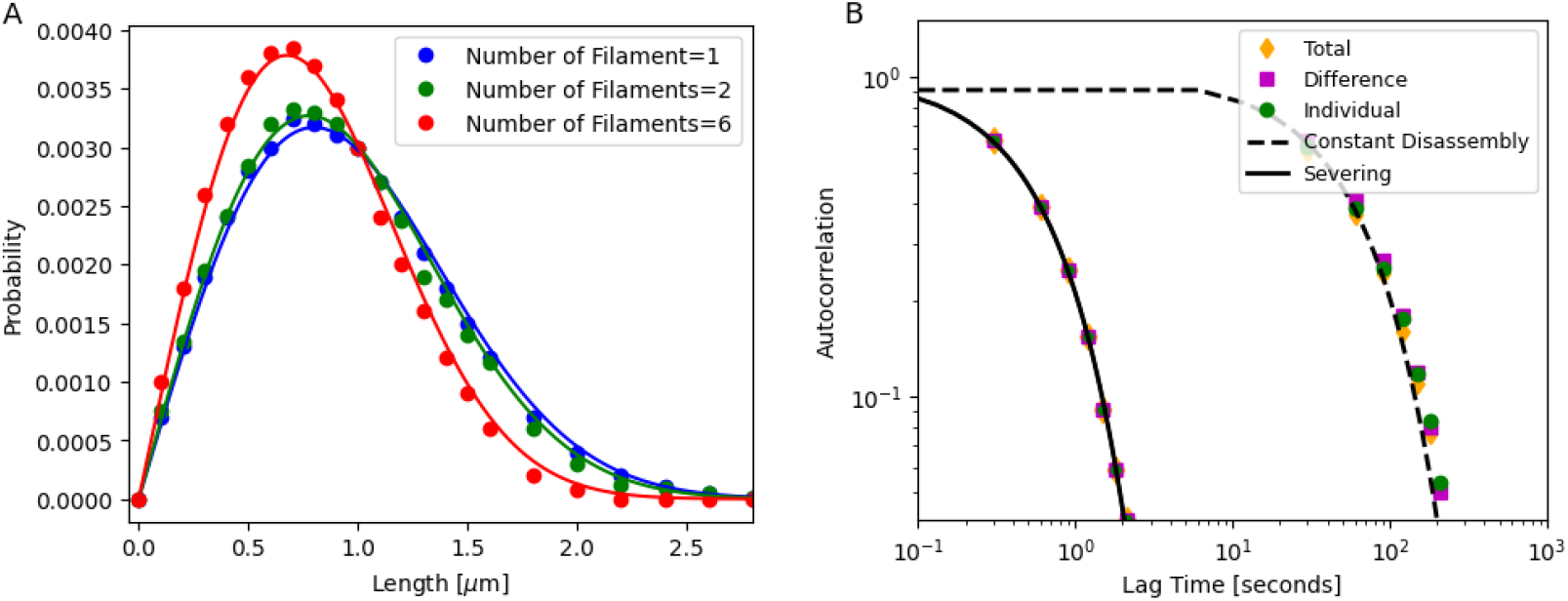
Length distribution and autocorrelation analysis of multiple filaments and bundles under various assembly-disassembly mechanisms. (A) Steady-state length distributions for *n* = 1, 2, 6 filaments regulated by severing in limited pool. Dots represent simulation results, while lines indicate analytical predictions (see Supplementary Material). Parameters: *N* = 3000 (monomers, each 4 nm in size), 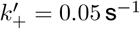, and *s* = 0.00375 monomers^−1^ s^−1^ (B) Autocorrelation functions for two co-assembling bundles in limited pool under constant disassembly (dashed lines) and severing (solid lines) mechanisms, showing individual filaments, their sum, and their difference. The plots are shown on a log-log scale, with dots representing simulation results, black solid and dashed lines indicating analytical predictions and fitted data, respectively (see Supplementary Material). Parameters: each bundle contains 4 filaments, with *N* = 100,000, 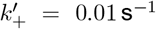, *k*_−_ = 1008.2 s^−1^ (constant disassembly), and *s* = 0.062 monomer^−1^ s^−1^ (severing).

**Figure S6.**
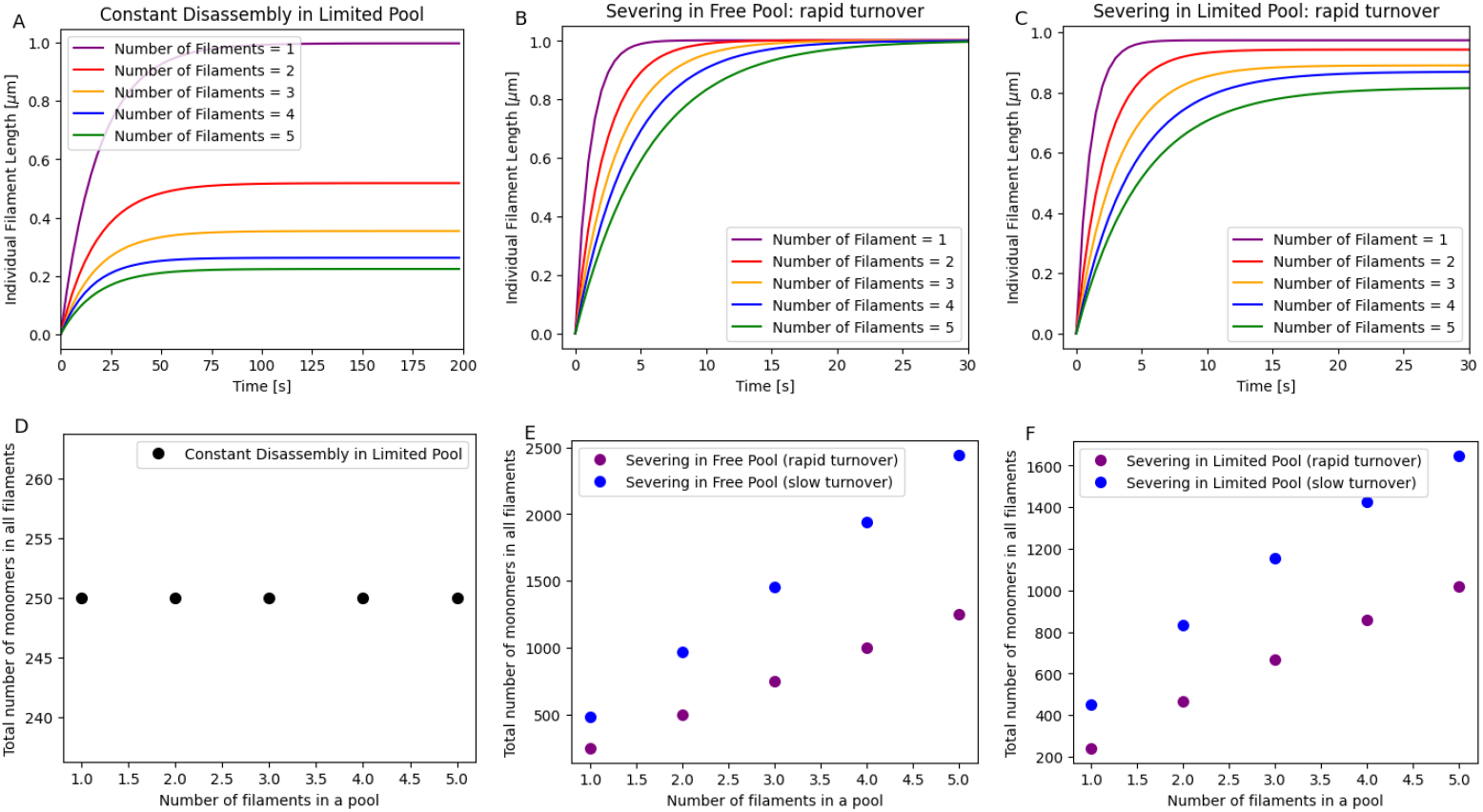
Assembly dynamics of multiple filaments under different conditions: (1) constant disassembly in a limited pool, (2) severing in a free pool, and (3) severing in a limited pool. (A–C) Stochastic simulations of the mean length of multiple filaments over time for each mechanism. (D–F) Total number of monomers incorporated into all filaments for each mechanism. Parameters: *N* = 3000 monomers (each 4 nm in size), with 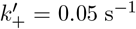 for all mechanisms. For the constant disassembly in a limited pool, *k*_−_ = 137.5 s^−1^; for the severing in a free pool and the severing in a limited pool, severing rates were *s* = 0.00375 monomers^−1^ s^−1^ (rapid turnover) and *s* = 0.001 monomers^−1^ s^−1^ (slow turnover).

**Figure S7.**
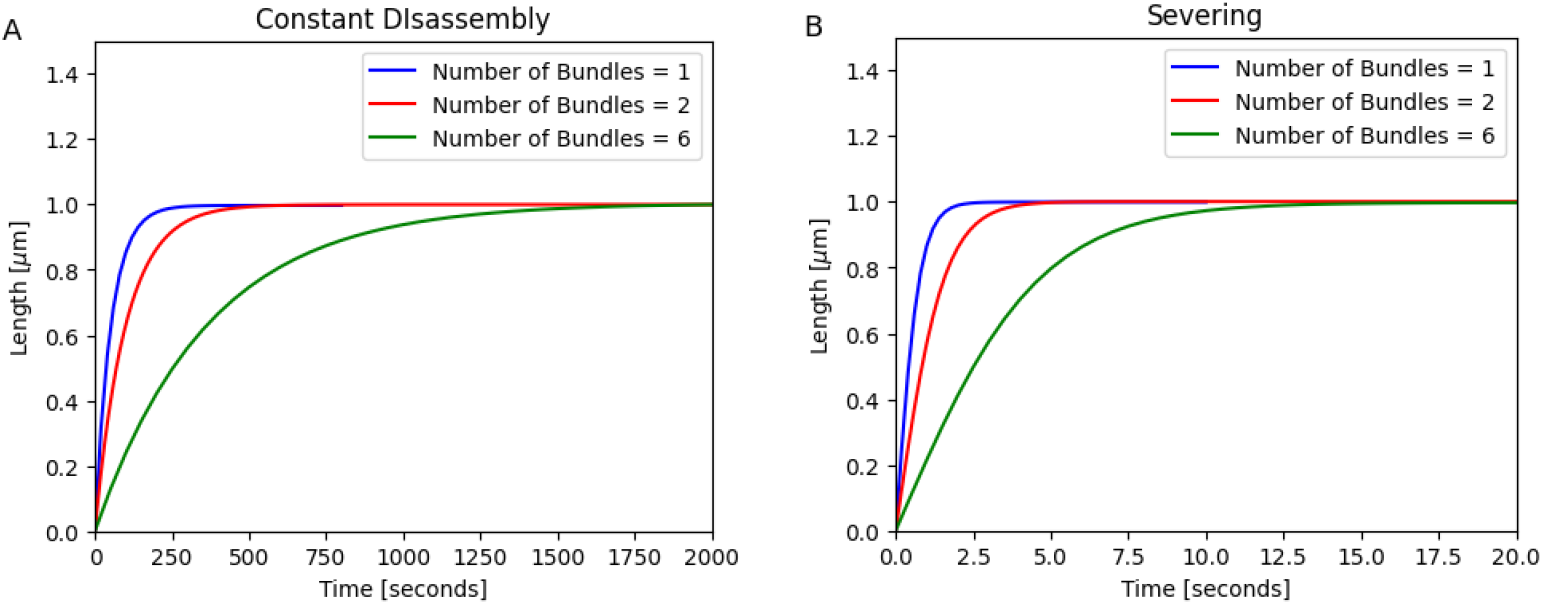
Growth dynamics of multiple bundles in a free pool with (1) constant disassembly and (2) severing mechanisms. (A–B) Stochastic simulations of the mean bundle length over time for both mechanisms. Parameters: each bundle contains 4 filaments, with *N* = 100,000 monomers (each 4 nm in size), and 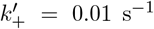. For constant disassembly, *k*_−_ = 1008.2 s^−1^; for the severing, *s* = 0.062 monomers^−1^ s^−1^.

